# Acoustic pH sensor for dynamic ultrasound imaging of cellular acidification

**DOI:** 10.1101/2025.01.24.634762

**Authors:** Dion Terwiel, Byung Min Park, Baptiste Heiles, Rick Waasdorp, Eleonora Muñoz-Ibarra, Tarannum Ara, Valeria Gazzola, David Maresca

**Affiliations:** Department of Imaging Physics, Delft University of Technology, Delft, the Netherlands; Netherlands Institute for Neuroscience, Royal Netherlands Academy of Art and Science, Amsterdam, the Netherlands

**Keywords:** biomolecular ultrasound, pH sensor, gas vesicles, lysosomal acidification, cross amplitude modulation

## Abstract

Genetically encoded pH sensors based on fluorescent proteins enable dynamic optical imaging of cellular processes such as endocytosis and exocytosis. To date, light scattering in thick tissue as well as photobleaching of fluorescent proteins prevent deep cellular imaging over sustained periods of time. To visualize intracellular pH variations across opaque organs, we introduce a genetically encoded acoustic pH sensor dubbed ‘pHonon’. We modified the outer gas vesicle protein (GvpC) of echogenic protein nanostructures via histidine point mutations. At low pH, engineered gas vesicles exhibit an increased shell stiffness which switched their acoustic response from nonlinear to linear. By combining pHonons with nonlinear ultrasound imaging, we captured dynamic deep tissue images of lysosomal acidification by macrophages in murine liver. The combination of pHonon with nonlinear ultrasound creates the possibility for basic studies of endo- and exocytic activity in deep tissue of living opaque organisms.

## 1 Introduction

pH regulation plays a crucial role in maintaining the homeostasis of cellular environments in healthy organisms. Disruptions of the pH balance are associated with numerous pathologies, including cancer ^1,2^, cardio- and cerebrovascular damage ^3^ or ischemia-reperfusion injury ^4^. The development of methods to visualize intra-cellular or extracellular pH variations in living tissue is therefore an active field of research ^5–9^. Because of light scattering in tick tissue, optical pH imaging methods are limited to studying thin specimens and cannot easily monitor pH variations at the organ scale. On the other hand, magnetic resonance pH imaging is sensitive to extracellular pH levels but cannot observe intracellular events.

In comparison, ultrasound provides an interesting trade-off in terms of spatiotemporal resolution, coverage, scalability, and availability of genetically encoded probes ^10,11^. Acoustic waves in the MHz range travel centimeters deep within opaque tissue while providing a resolution ranging from approximately 1 mm to 100 µm. Another advantage is that ultrasound imaging is performed with a single lightweight probe for transmission/reception, which provides easier access to specimens than microscopes or MRI scanners. Finally, the recent development of acoustic reporter genes and biosensors based on genetically encoded gas vesicles (GVs) ^12,13^ redefines the ability of ultrasound to visualize cellular function.

Assembled GVs consist of two structural proteins, gas vesicle protein A (GvpA) monomers and gas vesicle protein C (GvpC) ^14–16^. GvpA forms the outer wall of GVs in a helical chain, while GvpC provides mechanical reinforcement and rigidity to the shell. GvpC is currently the focal point for engineering GV acoustic sensors ^17,18^. Previous studies have shown that removal of GvpC results in a weakening of the GV shell ^19,20^, which allows GV shell buckling under acoustic pressure and triggers nonlinear ultrasound backscattering ^21^. The recent development of the first GV-based biosensor showed that the inclusion of protease recognition motifs in GvpC allowed proteases to cleave GvpC, thereby creating an ultrasound sensor that switched its acoustic response from linear to nonlinear in the presence of enzyme activity ^17^. More recently, a first ultrasonic calcium indicator based on GVs was reported ^18^. This sensor, which borrows design elements of fluorescent calcium indicators, relied on the inclusion of calmodulin in GvpC and of a calmodulin binding peptide linked through a flexible linker to the C-terminus of GvpC ^18^. In practice, binding of the calmodulin binding peptide to calmodulin and the subsequent deformation of GvpC causes GvpC to detach from GvpA, leading to a nonlinear acoustic response of the indicator in the presence of calcium.

Here, we present the development of the first GV-based acoustic pH biosensor for ultrasound imaging of intracellular acidification, dubbed ‘pHonon’. Leveraging insights from pH-sensitive fluorescent proteins as pHluorins ^5^ and our own analysis of the Cryo-EM structure of GVs ^15^, we inserted pH-sensitive amino acids into the GvpC sequence. Histidine is a polar amino acid that is often involved in pH-dependent protein activity in nature ^22–24^. The reason is that the chemical properties of the protonated and deprotonated histidine side chains are very different. At neutral pH the His side chain is uncharged, hydrophilic and apolar, while at low pH (*<*6) the side chain is positively charged, hydrophilic and polar. By replacing amino acids with histidine on the non-binding side of GvpC using structural insights from Cryo-EM ^15,16^, we aimed to trigger an allosteric conformational change in GvpC that would alter the GV wall stiffness and, eventually, modulate nonlinear ultrasound scattering of engineered GVs in response to a change in pH.

We designed and screened a small library of rationally engineered GvpCs with histidine point mutations and show that one of these mutants exhibits a pH-sensitive behavior under hydrostatic pressure and non-linear ultrasound imaging. We characterized this sensor dubbed ‘pHonon’ to unravel its presumed mechanistic behavior, report its kinetics, and present its performance in combination with nonlinear ultrasound imaging. To demonstrate the in vivo potential of pHonon biosensors, we performed ultrasound imaging of phagolysosomal acidification in liver macrophages ^25^. We show that pHonons are responsive to lysosomal acidification and report these changes in nonlinear ultrasound images. The parallel development of pHonon biosensors and volumetric imaging methods such as nonlinear sound-sheet microscopy ^26^ paves the way for deep and dynamic imaging of intracellular acidification in health and disease.

## 2 Results

### 2.1 pH-sensitive acoustic biosensor design and screening

To achieve a pH-dependent modulation of GV stiffness, we engineered the reinforcing protein GvpC with histidine substitutions on the solvent-facing side (**Fig. 1a**). A recombinant GvpC gene was expressed in *Escherichia coli* (*E. coli*), while GVs were purified from their native species (*Dolichospermum flos-aquae*) and stripped of GvpC. The GvpA shell and heterologously expressed GvpC were reconstituted in vitro (**Fig. 1b**) following a GV engineered pipeline developed by Lakshmanan et. al. ^20^.

**Figure 1.**
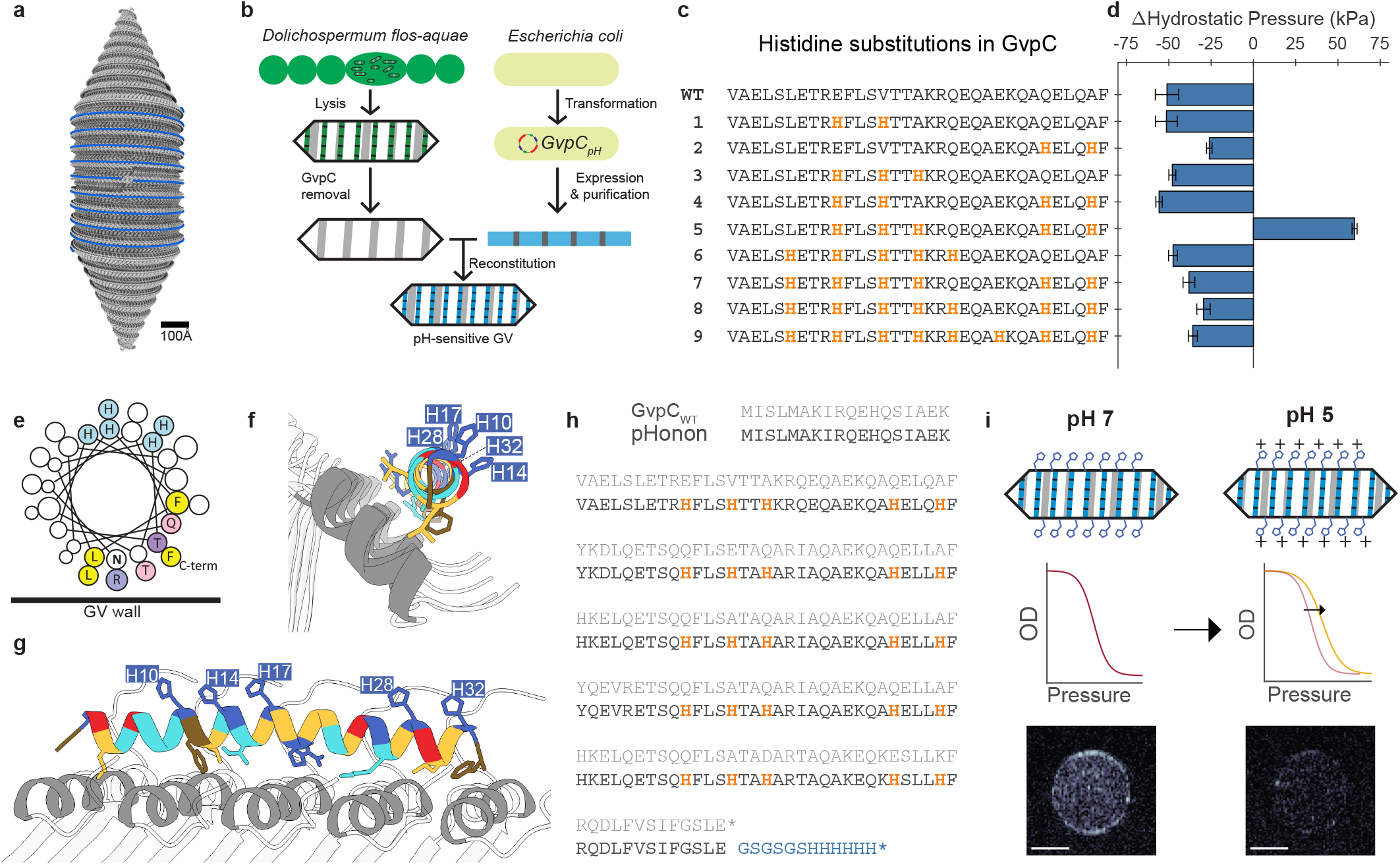
pH-sensitive GV design and screening. **a**, Molecular model of a GV; Grey: GvpA. Blue: GvpC. (adapted with permission ^15^). **b**, Schematic representation of production and reconstitution of recombinant GvpC on a stripped GvpA shell from *D. flos-aquae*. **c**, List of GvpC variants; only the amino acid sequence of the first repeat of GvpC is shown. Histidine mutations are shown in yellow. **d**, Change in the midpoint of hydrostatic collapse pressure between pH 7 (n=3) and pH 5 (n=3) for each GV variant. **e**, Helical wheel representation of variant 5, further referred to as ‘pHonon’. **f**,**g**, a 3D representation of a single repeat of pHonon bound to the GV wall (**f**, longitudinal plane; **g**, transverse plane). Amino acids are colored according to physico-chemical properties; Blue: Lys, Arg, His. Yellow: Met, Ile, Leu, Ala, Val. Light blue: Ser, Thr, Gln, Asn. Brown: Phe, Trp, Tyr. Red: Glu, Asp. **h**, Full amino acid sequence of wildtype *D. flos-aquae* GvpC (GvpC_*W T*_) and pHonon GvpC. Yellow: histidine mutations. Blue: Gly-Ser linker and His-tag. **i**, Schematic overview of pHonon function: low pH results in a positive charge on His residues. The resulting change in GvpC binding and conformation leads to a stiffer GV. The increase in stiffness reduces the contrast in non-linear ultrasound images.

We assembled a small library of 9 GvpC variants with different histidine substitutions (**Fig. 1c**). The locations of the histidine substitutions were chosen to be on the solvent-facing side of the GvpC, and determined by computational models of GvpC docking to GvpA ^15^. The position of each substitution was conserved with respect to the start of each repeat, meaning that each histidine occupies roughly the same position in each repeat regardless of the original amino acids’ physicochemical properties. Among the 9 different variants, only variant 5, dubbed ‘pHonon,’ exhibited an increased hydrostatic collapse pressure post-acidification (**Fig. 1d, Fig. A1**). pHonon has a total of 25 histidine substitutions, five in each repeat: H10, H14, H17, H28, and H32 (**Fig. 1e-h**). A conceptual sketch of the acoustic pH sensor behavior is provided in **Fig. 1i**.

### 2.2 Characterization and mechanistic studies of pHonon

In order to facilitate further engineering of pHonon biosensors, we explored the mechanism conferring pH-dependent GV shell strengthening. Contrary to wild-type and GVs without GvpC (ΔGvpC), pHonon biosen-sors exhibit an increased collapse pressure mid-point *Cp*_1*/*2_ at pH 5(**Fig. 2a-c**)^15^. pHonon shows a continuous strengthening over a time span of ∼20 hours, but 30% of the change in *Cp*_1*/*2_ occurs during the first 30 minutes after exposure to pH 5 (**Fig. 2d**). Experimentally, we observed batch variability with regards to pHonon *Cp*_1*/*2_ values at 20 hours, with one outlier that reached a Δ*Cp*_1*/*2_ of 150 kPa. Despite positioning all histidine substitutions on the solvent-facing side of GvpC binding residues, pHonons consistently showed a lower collapse pressure at pH 7 than wildtype GVs and GVs reconstituted with his-tagged GvpC.

**Figure 2.**
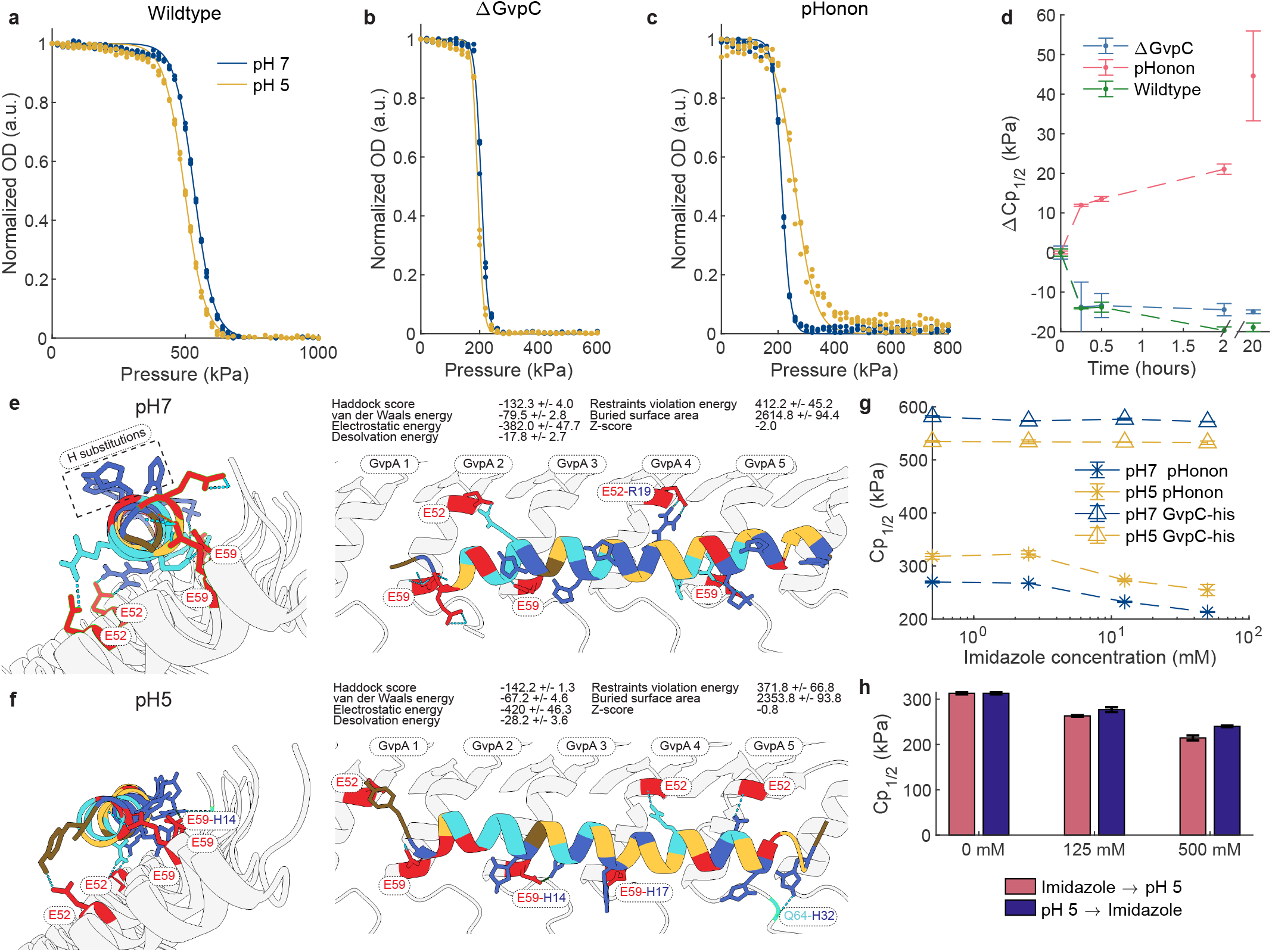
pHonon characterization and mechanism. **a-c**, Hydrostatic collapse pressure of wildtype GVs, ΔGvpC GVs and pHonon at pH7 (blue) and pH5 (yellow) (n = 3). The line is a sigmoidal fit. **d**, Time course of the hydrostatic collapse midpoint (*Cp*_1*/*2_) of wildtype GVs (green), ΔGvpC GVs (blue) and pHonon (red). *Cp*_1*/*2_ was determined using a weighted sigmoidal fit.**e**, Computational docking model of pHonon at pH7 generated using HADDOCK 2.4. **f**, Computational docking model of pHonon at pH5 generated using HADDOCK 2.4 **g**, *Cp*_1*/*2_ for different concentrations of imidazole present during the reconstitution of GvpA and GvpC. **h**, Imidazole competition assay. The effect of imidazole competing with His substitutions in GvpC for binding- and interaction sites on the surface of GvpA is shown by comparing the effect of adding the competitor (imidazole) before the hypothetical allosteric conformation change (red) or after (blue).

We used computational docking ^27,28^ to gain insights into possible binding orientations of pHonon GvpC with GvpA at physiological and acidic pH (**Fig. 2e,f**). To limit the computational complexity of the docking we used Cryo-EM data ^15^ to constrain the GvpA residues involved in binding to alpha-helix 2 (I37 to I46), and the GvpC binding residues to residues 4, 10, 11, 12, 14, 15, 17, 19, 23, 26, 28, 30, 32, 33 (highly conserved residues plus the newly introduced histidine mutations). At physiological pH, the highest scoring cluster (**Fig. 2e**) has a very similar orientation to wildtype GvpC, as previously reported ^15^. A higher docking score was observed for pHonon, which we primarily attribute to a slight tilt of pHonon GvpC, allowing E59 on GvpA2 to form hydrogen bridges with K2 on GvpC. Histidine point mutations point outward in relation to the GV wall as anticipated.

Acidic pH was simulated by manually changing histidine protonation states to fully protonated, i.e., with a positively charged side-chain. The highest score for pHonon at pH 5 shows GvpC rotated along its long axis (**Fig. 2f**). Histidine substitutions now point parallel or inwards towards GvpA alpha-helix 2. Salt-bridges are predicted between E59 on GvpA2 and GvpA3 with H14 and H17 respectively. Hydrogen bridges are also predicted between GvpA1:E52-F1, GvpA2:E59-K3, GvpA4:E52-Q22 and GvpA5:E52-K25. The hydrogen bridge GvpA5:Q64-H32 may be inaccurate as the N-terminal tail is unfolded and can take any orientation or conformation, but was here modeled as a rigid structure. A second high scoring structure was found with a similar orientation at pH 7. Follow-up studies should investigate whether both structures exist and are in competition, or whether only one of the two occurs experimentally.

To determine if the docking model at pH 5 (**Fig. 2f**) is plausible, we performed two competition assays with the histidine analog imidazole to investigate interactions between histidine and GvpA. According to the most likely docking, adding imidazole at pH 7 should not interfere with any bond formation as none of the histidine substitutions show an interaction.

In the first competition assay (**Fig. 2g**), we added imidazole during the reconstitution of GvpC-his and pHonon respectively. Increased concentrations of imidazole during the reconstitution process led to lower pHonon collapse pressure by 40 kPa (**Fig. 2g**). Collapse pressures at pH 5 were measured to be consistently 40 kPa higher than the collapse pressure at pH 7. Imidazole had no effect on GvpC-his reconstitution. Based on the docking results, we did not expect imidazole to affect pHonon reconstitution. However, solubility might be a confounding factor in these measurements as solubility of GvpC-his at pH 7 is significantly higher than that of pHonon (**Fig.A2**). We suspect that a considerable amount of pHonon GvpC may precipitate out of solution as urea is removed before it gets to bind GvpA. Interestingly, this result shows that we can control the initial collapse pressure of pHonon at pH 7 and therefore modulate the initial level of nonlinear scattering that a pHonon biosensor would have. Preparing pHonon sensors with different initial collapse pressures would enable multiplexing ^29^.

The second competition assay aimed to investigate whether the rotated GvpC docking is realistic (**Fig. 2h**). If pHonon GvpC truly rotates at acidic pH, imidazole should compete with the newly formed salt-bridges (E59-H14, E59-H17) and diminish the characteristic strengthening of pHonon. Imidazole was added either before or after incubation at pH 5 for 20 hours, and the collapse pressure was evaluated. Adding imidazole before changing the pH resulted in a lower collapse pressure than adding imidazole after changing the pH (**Fig. 2h**). We interpret this result as imidazole partially blocking the formation of salt-bridges (E59-H14, E59-H17), leading to a lower collapse pressure. When imidazole was added after changing the pH, the already formed bond blocked an interaction with imidazole and the resulting GV had a higher collapse pressure. While these computational docking results produce intriguing and stable structures, it is important to note that they remain largely speculative. For example, the assumption of a uniform pH - where all amino acids experience the same pH level - is almost certainly inaccurate. The biochemical microenvironment of each amino acid likely exposes it to significantly different pH levels. Furthermore, the assumption that a higher HADDOCK score corresponds to a stiffer GV may not hold true, as higher association energies do not necessarily imply changes in the protein’s mechanical properties. Nevertheless, the proposed dockings provide a robust starting point for a more detailed exploration of the molecular mechanisms underlying pHonon.

### 2.3 Dynamic ultrasound imaging of acidification

We performed ultrasound imaging of pHonon and control nanoparticles (ΔGvpC GVs) embedded in 2% agarose phantoms using a nonlinear cross amplitude modulation pulse sequence (xAM) ^21^ (**Fig. 3a**).

**Figure 3.**
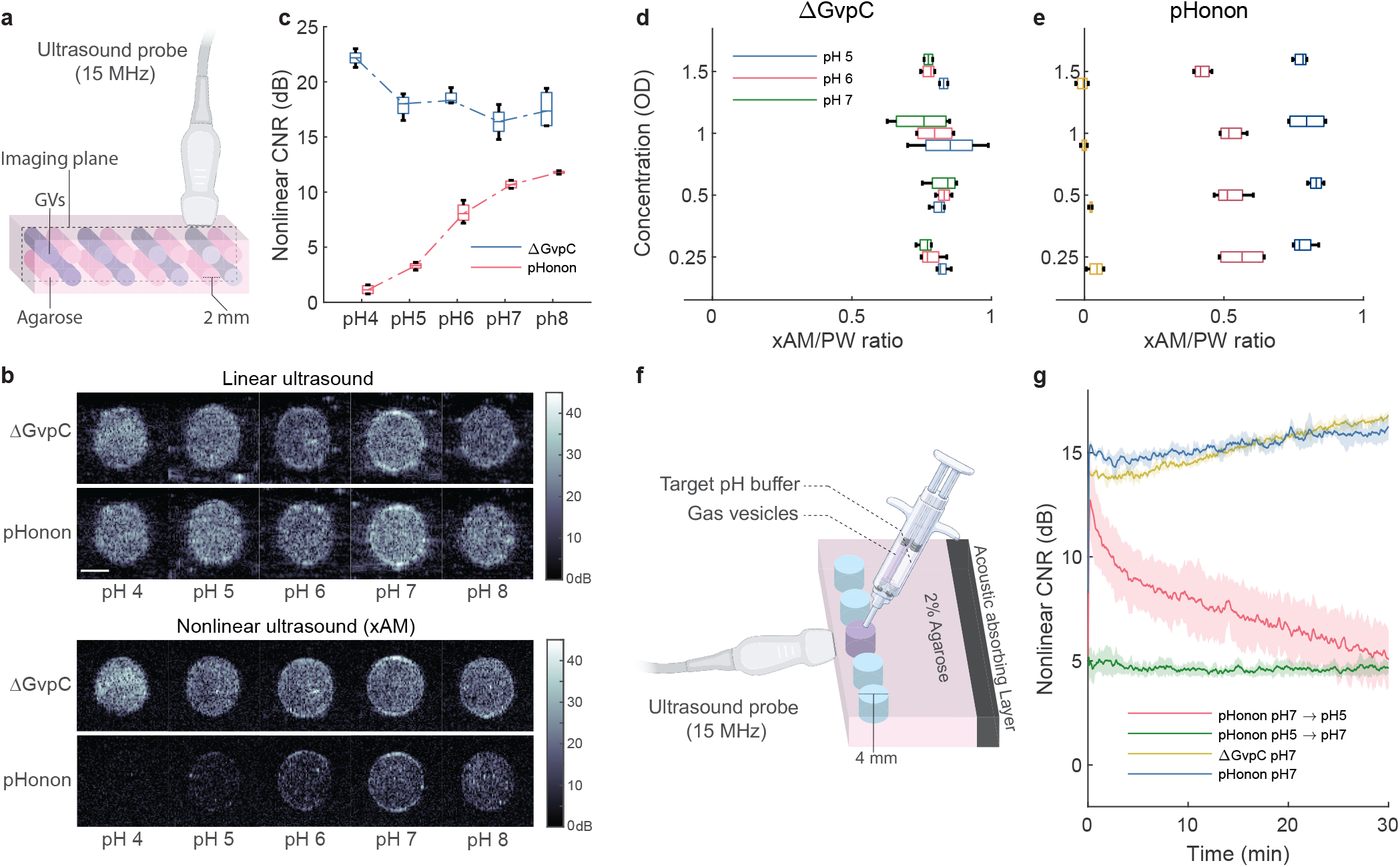
In vitro ultrasound characterization of pHonon. **a**, Schematic view of GV phantom imaging. **b**, Linear (top) and nonlinear (bottom) ultrasound images of Δ GvpC and pHonon GVs (OD = 1) after overnight incubation at the target pH and immobilized in 2% agarose. **c**, Nonlinear contrast-to-noise ratio (CNR) of ΔGvpC (blue) and pHonon (red) (n = 4). The region of interest(ROI) was selected over the entire well, and the noise level was determined using an equally sized ROI over a well filled with 2% agarose. **d**, The ratio of nonlinear (xAM)/linear (plane wave(PW)) CNR of phantom images of ΔGvpC and pHonon (n = 4) (**e**) for a range of pH values from 5 to 7 and concentrations from OD 0.25 to OD = 1.5. **f**, A schematic view of the kinetics measurement setup where a 4mm well in 2% agarose is filled from a dual-barrel syringe with equal volume OD = 1.5 GVs and 0.3 M McIlvaine buffer at the target pH. Imaging is performed at a frame rate of ∼2 Hz and the probe is coupled with ultrasound gel. Blue wells are filled with demiwater to minimize reflection artifacts **g**, Nonlinear CNR over time with the injection and mixing of GVs with a target pH buffer at time 0 . pHonon is exposed to pH 5 (red) and subsequently to pH 7 (green). pHonon and ΔGvpC at constant pH (blue,yellow) were used as controls. (n = 5)

Linear ultrasound images revealed a similar echogenicity for both ΔGvpC GVs and pHonon across the pH 8 to 4 range (**Fig. 3b**). Under nonlinear ultrasound exposure, we observed a 10.6 *±* 0.4 dB decrease in the non-linear contrast-to-noise ratio (CNR) of pHonon at pH 4 (**Fig. 3c**) whereas control GVs remained echogenic through the pH range explored. The ecliptic behavior of engineered pHonon nanoparticles is unique and aligns with findings from collapse pressure measurements reported in (**Fig. 2**). We also measured an increase of the CNR of ΔGvpC GVs of 4.8 ± 1.3 dB at pH 4. This drop in CNR coincides with a weakening of the pHonon nanoparticles observed in hydrostatic collapse pressure measurements (**Fig. 2a**). Development of a quantitative ultrasound readout of pH levels in tissue is critical if we are to use pHonon for deep tissue imaging. Two physical effects in particular can lead to confounding effects: depth-dependent ultrasound attenuation in tissue and local variations in GV concentration. We developed a ratiometric measure, the cross amplitude over plane wave ratio (xAM/PW), that normalizes nonlinear scattering of GVs by their linear scattering which is proportional to GV concentration (**Fig. 3d-e**). The xAM/PW ratios of control ΔGvpC GVs was constant for any GV concentration (OD 0.25 to 1.5) or pH (5 to 7) tested, as expected from a nanoparticle which is not pH sensitive (**Fig. 3d**). For pHonon, xAM/PW ratios were independent of concentration but dependent on pH, with ratios varying from 0.80 at pH 7, 0.51 at pH 6, and 0.02 at pH 5 (**Fig. 3e**).

We measured the switching kinetics of pHonon using a custom-built experimental setup (**Fig. 3f**). In our experimental configuration, a 2% agarose phantom with 4 mm diameter open wells was molded as an acoustically transparent propagation medium. A 2x 1mL dual-barrel syringe allowed the rapid mixing of 0.3 M pH 5 McIlvaine buffer with GVs at OD 3 in 0.05% PBS, followed by the direct injection of this mixture into a well. Ultrasound imaging of the mixed solution was conducted at a 2 Hz framerate over 30 minutes to monitor changes in nonlinear contrast caused by the rapid exposure to a target pH value (**Fig. 3g**). Upon quasi-instantaneous exposure to pH 5, the xAM CNR of pHonon decreased by 2.2 ± 1.1 dBs in 90s, 4.7 *±* 0.3 dBs in 10 min and 6.1 ± 0.85 dBs in 20 min. We assessed the reversibility of our pH-sensitive GVs by exposing pHonon incubated at pH5 for 20 minutes to a quasi-instantaneous pH 7 buffer. We observed that xAM CNR remained constant at its pH5 value over the course of 30 minutes, indicating that our sensor design is not reversible between pH 7 and pH 5. ΔGvpC controls did not show any pH dependence as expected from imaging results.

### 2.4 *In vivo* ultrasound imaging of lysosomal acidification in liver macrophages

After characterizing pHonon in vitro, we assessed the ability of pHonon biosensors to report on intracellular acidification in living tissue. We chose to perform dynamic imaging of phagolysosomal acidification in liver macrophages, as macrophage uptake of intravenously administered GVs is well documented ^12,25^. Kupffer cells, the liver’s primary phagocytic macrophages, are primarily found in the sinusoidal spaces of the liver ^30^. Foreign particles such as GVs are captured on the surface of Kupffer cells and subsequently phagocytosed. Next, the phagosome, now containing GVs, merges with lysosomes and gets acidified down to a pH of 4.5^31^. It is acknowledged that the pH within phagolysosomes drops from 7 to approximately 5 over the course of 10 to 20 minutes ^32,33^. Previously reported biomolecular ultrasound experiments also show that GVs circulating in the blood stream primarily accumulate in the liver ^25^. Following intravenous injection, ultrasound contrast generated by GVs builds up in the liver over the course of 10 minutes before nearly disappearing after 60 minutes ^25^.

A jugular vein catheter was placed in anesthetized C57BL/6 mice. After preparation of the animal, the left lateral lobe was found by using ultrafast Doppler. When possible, the caudate lobe was left out of the imaging plane. An xAM imaging sequence operating at a framerate of 2 Hz was used to record data for 45 minutes. After two minutes, the animal received a 150 *µ*L bolus of either pHonon or control ΔGvpC GVs at OD = 50 through the catheter (**Fig. 4a**). The catheter was then flushed with saline (see Methods).

**Figure 4.**
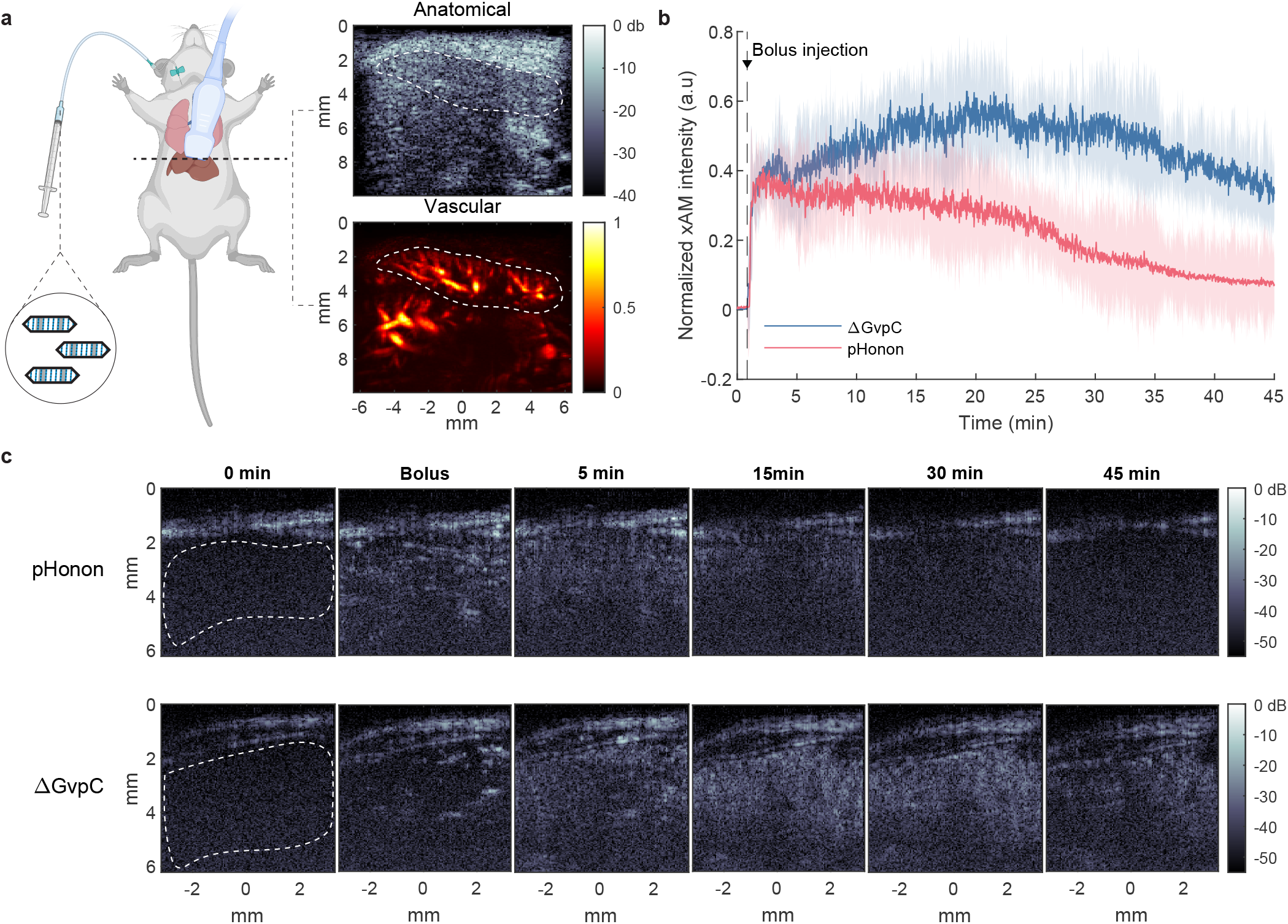
In vivo nonlinear imaging of lysosomal acidification in mice livers. **a**, Schematic showing the experimental setup. Shown top right is a representative anatomical (planewave) ultrasound image, and lower right a vascular (doppler) ultrasound image of a mouse liver. **b**, Average normalized nonlinear (xAM) signal intensity in the liver after bolus injection of ΔGvpC (blue) or pHonon (red) GVs (n = 4). **c**, Representative nonlinear ultrasound images of ΔGvpC (bottom) or pHonon (top) GVs in murine livers of two different animals at different time points. The ROI for these specific mice is indicated at t = 0 minutes with a white dotted line.

For ΔGvpC GVs, we measured a rapid increase in xAM signal due to the bolus injection and the intravascular circulation of GVs in liver vessels. This was immediately followed by a slower increase in xAM signal in the liver plane of interest over 20 minutes post-bolus, and followed by a slow decrease until the end of the 45 minutes recordings (**Fig. 4b,c**). This observation is in agreement with the literature ^25^.

For pHonon on the other hand, after the initial increase in xAM signal due to the bolus injection and the intravascular circulation of pHonons in liver vessels, we measured a steady decline of the xAM signal in the liver plane of interest until the end of the 45 minutes recordings. At the peak of the bolus, a “spotty” xAM signal was observed which we attributed to perfusion of the sinusoidal spaces. At this stage, GVs are thought to be confined in the vascular space rather than internalized in liver macrophages (**Fig. 4c**). Over the next 5 minutes the xAM signal spread out through the plane as GVs get internalized into phagosomes. Phagosomal acidification is a biological process that takes place during and shortly after the initial lysosomal merging. In our experiments, pHonon biosensors clearly displayed a lower level of xAM signal compared to ΔGvpC GVs as of 15 mins.

Note that for ΔGvpC GVs, the loss of xAM signal after 20 minutes is caused by proteolytic degradation of the GVs rather than acidification. Together, our results indicate that phagolysosomal acidification takes place within the initial 20 minutes of the experiment, which is only slightly longer than the literature reports of acidification times of 17 min, 15 min, and 10 min respectively ^31–33^. Altough it should be remarked that most of these studies found large variations in overal acidification times as well. This result is also consistent with the ecliptic behaviour of the sensor reported in **Fig. 3**. Representative xAM images at different time points of the recording are reported in **Fig. 4c**.

## 3 Discussion

We successfully engineered an acoustic pH biosensor capable of reporting lysosomal acidification in ultrasound images. Our approach consisted of assembling a small library of nine recombinant GvpCs containing histidine point mutations, because this amino acid has the desired pKa for a sensor of endocytosis. We introduced histidine at positions that do not interfere with binding of GvpC to GvpA and screened for engineered GV variants that would show an increased shell stiffness when exposed to a target pH of 5, which is the acidic value reached in late phagolysosomes ^31–34^. One of the engineered GV variants dubbed pHonon showed a 45 kPa increase in hydrostatic collapse pressure at low pH and was selected for further analysis. All other variants of the library showed weakening at low pH. Computational modeling provided insights into the potential sensing mechanism of our pH sensor. Protein docking scores indicated that at pH 7, pHonon GvpC residues form hydrogen bridges with GvpA . At pH 5, pHonon GvpC rotated along its axis to form salt bridges and hydrogen bridges with GvpA, further strengthening the shell. Interestingly, the addition of a histidine analog during pHonon reconstitution conferred a handle to modulate the biosensor shell stiffness at pH 7. Because the collapse pressure mid-point is directly related to the level of nonlinear scattering at pH 7, multiplexed biomolecular ultrasound imaging ^29^ could be investigated in the future. pHonon biosensors successfully reported pH variations in ultrasound images, exhibiting a 10.6 dB dynamic range between nonlinear ultrasound contrast at pH 7 and pH 4. In terms of kinetics, a 6 dB drop in nonlinear ultrasound contrast was measured in 20 min. This ecliptic sensor behavior is similar to pH-sensitive fluorescent proteins developed by Miesenbock et. al ^5^. Normalizing the nonlinear acoustic response of pHonon with their linear acoustic response enabled ratiometric imaging of pHonon biosensors at pH 7, pH 6 and pH 5 respectively. Future work will evaluate this approach in an in vivo context to test the feasibility of parametric ultrasound mapping of pH levels in tissue. We demonstrated the potential of acoustic pH biosensors by imaging lysosomal acidification in liver macrophages. Following intravenous injection, the amplitude of nonlinear pHonon signals in the liver decreased steadily because of internalization in macrophages and exposure to low pH conditions. Control GVs showed a steady increase in nonlinear signal until 20 minutes before decreasing due to enzymatic protein degradation.

While we succeeded in engineering a pH-sensitive GV and imaged cellular acidification in vivo, pHonon should be seen as a first step towards a class of pH-sensitive GV-based biosensors. Our semi-rational engineering will serve as a basis for further optimization and the development of acoustic pH sensors with enhanced sensitivity, different pH transitions, potentially faster kinetics or reversibility, similarly to the successive improvements of optical pH sensors similarly to the successive improvements of optical pH sensors ^5,6,34–37^. Interestingly, pHonon is the first GV-based sensor to show a shell strengthening, as opposed to a shell weakening seen in other GV-based biosensors ^17,18^, providing a new mechanistic cue for GV engineering. While our current approach relied on a limited screening library, the recent development of high-throughput directed evolution assays for GV engineering ^38^ could expand this variant-space.

The exact pH sensing mechanism of pHonon biosensors remains to be elucidated. The present study reports plausible mechanisms based on a computational docking model and limited experimental data. Several questions remain open. For example, it would be interesting to investigate why structurally similar GvpC mutants of our library did not lead to pH-dependent shell strengthening. Current docking models seem to point at H14 and H17 as key positions, which is a combination of mutations that also occurred in variants 3 and 6-9 of our library. A combination of further mutagenic study, combined with computational models and high-resolution microscopy techniques might provide novel insights.

The clinical relevance of these pHonon biosensors was not the aim of this study but it will be interesting to revisit molecular ultrasound imaging methods such as ischemic memory imaging ^4^ and to explore new avenues for molecular ultrasound imaging such as monitoring of impaired lysosomal function in neurodegenerative diseases ^39^.

In conclusion, our results demonstrate that protein-based acoustic pH sensors can enable dynamic ultrasound imaging of intracellular acidification in intact opaque organisms and paves the way for biomolecular ultrasound imaging of organelle function.

## 4 Methods

All chemicals were purchased from Merck (Sigma Aldrich) unless specified.

### 4.1 Gas vesicle production

*Dolichospermum flos-aquae* (CCAP 1403/13F; Culture Collection of Algae and Protozoa) was cultured according to previously published protocol ^40^. *D. flos-aquae* was cultivated for two weeks in G625 medium (5.84 mM NaNO_3_, 224 *µ*M KH_2_PO_4_, 304 *µ*M MgSO_3_·7H_2_O, 208 *µ*M Na_2_SiO_3_·9H_2_O, 189 *µ*M NaCO_3_, 10 mM NaHCO_3_, 245 *µ*M CaCl_3_, 31 *µ*M citric acid and 3 *µ*M EDTA) supplemented with BG-11 freshwater medium (Sigma Aldrich, C3061) at 25 ^*°*^C, 120 rpm and 1% CO_2_. The culture was illuminated at 5000 lux on a 14 h light, 10 h dark cycle. Cultures were allowed to settle for 24-48 hours, and the floating cell layer was collected. Harvested cells were lysed in the following buffer 10% v/v harvested cells, 10% v/v Solulyse^*T M*^ bacterial lysis agent (Genlantis, L200500), and 500mM sorbitol for 6-8h on a rotary mixer at room temperature. GVs were separated from the lysate by centrifugation at 350g for 12h. GVs are in the floating white layer, the subnatant was removed and discarded. The floating GV layer was topped up with phosphate buffered saline, pH 7.4 (PBS). The centrifugation and subnatant discard were repeated 4 times in total. GvpC was removed from GVs by 3 subsequent rounds of centrifugation at 350 g for 12 hours, subnatant discard and buffer top up, using PBS supplemented with 6 M urea as the buffer. GV concentration was measured by optical density measurements at 500 nm with a spectrophotometer (Ocean optics, STS-VIS).

### 4.2 Cloning methods

All gene sequences were codon-optimized for E. coli expression and inserted into pET28a plasmid via Gibson assembly. Enzymes were purchased from New England Biolabs and primers from Sigma Aldrich. The resulting plasmids were transformed into *E. coli* BL21(DE3) for expression.

### 4.3 Heterologous expression of GvpC in *E. coli*

Seed culture was prepared by inoculating LB with transformed colonies of BL21(DE3) and grown overnight at 37°C, 220 rpm. The overnight seed culture was inoculated into auto-induction Terrific Broth (TB). Auto-induction TB medium is composed of 24 g/L yeast extract, 12 g/L tryptone, 2.9 g/L lactose and 7.6 g/L glucose ^41^. The medium was sterilized via autoclave, and supplemented with 2% w/v lactose, 0.5% w/v glucose, and 100 *µ*g/L ampicillin. Cultures were collected and centrifuged at 3000 g for 15 mins. The resulting cell pellet was resuspended in bacterial lysis buffer (50 mM Tris-HCl pH 8, 100 mM NaCl, 1% v/v Triton X-100, 10 mg/mL lysozyme) and rotated for 1-2 hours at room temperature. The lysate was centrifuged at 18000 g for 40 minutes, and the supernatant was discarded. Inclusion body solubilization buffer (6 M Urea, 50 mM Tris, 100 mM NaCl) was used to resuspend the centrifuged pellet, and the resuspension was centrifuged at 18000 g for 40 minutes. The His-tagged GvpC proteins were purified using Ni-NTA affinity chromatography (HisPur, Thermo Fisher, 88221) following the manufacturer’s protocol. Protein concentration was determined using Bradford assay (Thermo Fisher, 23238) according to the manufacturer’s protocols, bovine serum albumin was used to make the standard curve.

### 4.4 Reconstitution of GV with GvpC

GvpC was re-added to ΔGvpC GVs in a 20x mole excess (GvpC:GvpA) according to the formula 20 × OD_500 nm_ × 198 nM × volume (in liters) of GVs = nmol of recombinant GvpC ^40^. The ΔGvpC GV and GvpC protein mixture was dialyzed twice against 2 L of PBS with a 6 kDa MWCO dialysis membrane. followed by 3 rounds of centrifugation at 350 g for 12 hours, subnatant discard and buffer top-up, using PBS as the buffer.

### 4.5 Hydrostatic collapse pressure measurements

Hydrostatic collapse pressures were determined using the setup outlined previously ^40^. GVs were diluted to an OD_500 nm_ of 0.2 with the appropriate buffer. McIlvaine buffer (citric acid, Na_2_HPO_4_) was used for all pH-dependent collapse pressure studies. A pressure controller (PC series; Alicat Scientific) connected to a 1.5 MPa nitrogen gas source was used to vary the hydrostatic pressure in a flow-through cuvette (Hellma, article no.1767001510-40). A pressure step of 20 kPa and an equilibration time of 5 seconds were used before the optical density at 500 nm was recorded using a spectrophotometer (STS-VIS; Ocean Optics). The pressure controller and spectrophotometer were controlled using MATLAB. All measures were done in triplicate and the average and 1 standard deviation are reported.

### 4.6 Computational docking

A docking model was generated using the online environment (https://rascar.science.uu.nl/haddock2.4/) for HADDOCK 2.4^27,28^. Previously published atomic models for docking of GvpC to GvpA ^15^ were retrieved (https://doi.org/10.5281/zenodo.6867443) and used as a starting point for our docking simulations. To transform GvpC from the published model to pHonon GvpC histidine mutations were programmed in ChimeraX ^42^ using the ‘swapaa’ command. Rotamers were left to their default settings. Active residues for GvpA were limited to residues 51–61 of GvpA, similar to previously published computational docking methods for GvpC ^15^. For GvpC, active residues were residues 4,10,11,12,14,15,17,19,23,26,28,30,32,33, and selected by adding the histidine mutations to the residue list used in a previously published docking ^15^. To simulate docking at pH 7 all HADDOCK settings were set to default. To simulate pH5 the histidine protonation states of GvpC were manually set to HIS+ in the HADDOCK environment. Docking clusters were displayed in ChimeraX and hydrogen bonds between GvpA and GvpC highlighted with the ‘hbonds’ command.

### 4.7 Competition assays

To investigate the effect of initial imidazole concentration during reconstitution, imidazole concentration in the reconstitution mixture was varied. Purified GvpC was subjected to several rounds of dialysis against 6M Urea and PBS to reduce the imidazole concentration to ¡ 10 *µ*M. Dialyzed and purified GvpC was mixed with imidazole to achieve reported final imidazole concentrations. This mixture was then combined with ΔGvpC GVs in a 20x mole excess and dialyzed for 2 rounds against 2 L of PBS.

pH-dependent competition of GvpC and imidazole was investigated by switching the order of the pH drop or imidazole addition of the pHonon GV solution. Imidazole was added to the pHonon GV solution to the reported final concentrations. The pH was dropped by adding 0.1M citric acid into the pHonon GV solution to reach pH 5. The GV solution was incubated for 15 minutes at each step, and the hydrostatic pressure was measured after both pH drop and imidazole addition.

### 4.8 In vitro Ultrasound imaging

All ultrasound imaging was performed with a Verasonics Vantage 256 scanner combined with an L22-14vX Verasonics ultrasound probe (Verasonics, Kirkland, Washington, USA), with a specified pitch of 0.1 mm, an elevation focus of 8 mm, an elevation aperture of 1.5 mm and a center frequency of 18.5 MHz. For the immobilized invitro GV images 2% agarose phantoms were prepared in Milli-Q or PBS using custom 3D-printed molds. GV samples were diluted 2x in 2% low melting-temperature (LMT) agarose and loaded into the phantom wells. GV samples were matched at OD_500 nm_ of 2. A well filled with 1% LMT agarose was used to determine background intensity in every frame. The cross-amplitude modulation (xAM) method for non-linear imaging of contrast agents was used ^21^. xBmode images were acquired using a single cross-propagating plane-wave transmission. Plane wave images were beamformed by coherent compounding of the two mirrored plane waves from the xAM pulse sequence. xAM images were acquired using three separate transmits, 2 mirrored plane waves and a cross propagating wave. Contrast-to-noise values were determined by taking the log-compressed ratio of the signal in a representative circular region of interest (ROI) in the well, and a representative noise ROI in a well filled with 1% LMT agarose. Images were displayed with an equalized noise-floor. All measures were done in quadruplicate and the average, 25% and 75% and minimum and maximum are reported.

To measure pH dependent kinetics of pHonons, a 2% agarose phantom with 4 mm wells was prepared. A dual-barrel syringe (2 x 1 mL) with a mixing tip was filled with GVs (OD_500 nm_ = 2, 0.5% PBS) in one barrel and 0.3 M McIlvaine buffer in the second barrel. Buffer capacity was calibrated to be pH 5 after injection into the well of the phantom. The ultrasound probe was positioned against the phantom and acoustically coupled using ultrasound gel. GVs and pH buffer were injected into a phantom well through the syringe mixing tip. Images were recorded at an acoustic pressure of 600 kPa, with a framerate of 2 Hz for 30 minutes.

### 4.9 In vivo imaging of phagolysosomal acidification

C57BL/6 mice of ages between 3 and 5 months were anesthetized with Isoflurane (5% for induction, 1% for maintenance) and placed on a heating pad with loop control from a temperature probe and breath monitoring. The animal was equipped with a foot sensor to measure pulse (MouseStat and Physio Suite from Kent Scientific, Torrington, Connecticut, USA). Two areas were shaved and depilated: the area over the right jugular vein to prepare for catheterization and a 4cm x 3cm area from the ribs down to allow for artefact-free imaging from the hair. The area over the right jugular vein was cleaned 3:3 times with saline solution and a mix of chlorhexidine/cetrimide (Hibicet, Molnlycke Health Care, Gothenburg, Sweden). After skin incision and resection of fatty tissue, the jugular vein was isolated and a catheter was placed inside and fixed with sterile non-resorbant sutures. The catheter was flushed with a mix of saline and heparin to prevent blood clots. Then the skin was sutured. The mice were hydrated during the imaging session through subcutaneous bolus of 0.1 mL of saline solution. Ultrasound gel was applied in the abdominal area as a coupling medium and an L22-14Vx ultrasound probe (as described in 4.8) was placed medial to the sternum. Through the use of ultrafast Doppler, the field of view was focused on the lateral left lobe and a plane containing a representative vascular tree was isolated. Non-linear xAM images were generated using 3 compounded angles (12^*°*^, 15^*°*^, 21^*°*^). A baseline ultrasound signal was established for 2 mins before a 150 µL bolus of GVs at OD_500 nm_ of 50 in PBS was administered through the vein catheter.

The catheter was flushed with saline to flush the dead volume in the catheter tubing. A maximum volume of 200 *µ*L was injected per experiment. Nonlinear images were acquired for 45 mins post-injection at a frame rate of 2 Hz.

Breathing creates a significant motion below the sternum, leading to motion artifacts. Since we are imaging in 2D, out-of-plane motion cannot be compensated for, and the imaging plane changes entirely during some periods of the breathing cycle. As our frame rate is high enough to image the relatively slow phagocytosis of the GVs and their pH-induced behavior changes, we elected to compensate for the motion by using a frame-rejection approach. The liver was manually segmented in the linear image extracted from the main pulse of the xAM and the average signal intensity within the ROI was calculated. A peak detection algorithm was used to find the signal intensities with the large deviations from the baseline. Frames were rejected at an intensity level of 50% of the average intensity of the detected peaks. To conserve uniform sampling of the nonlinear signal, rejected frame intensities were linearly interpolated from their previous and next samples. The resulting time-intensity curves were smoothed using a 5-point moving average window and normalized with respect to an average signal of the baseline set to zero and dividing by the signal maxima. Signal traces from each experimental group (pHonon n = 4, ΔGvpC n = 4) were averaged at each time point and plotted.

### 4.10 Key resource table

| Resource | Identifier |
| --- | --- |
| Haddock 2.4<sup>27,28</sup><br>ExPASy<sup>43</sup><br>ChimeraX 1.7<sup>42</sup> | https://wenmr.science.uu.nl/haddock2.4/<br>https://www.expasy.org/<br>https://www.rbvi.ucsf.edu/chimerax/ |
| Generated plasmids | Source |
| pET28a-GvpC-His6 | This study |
| pET28a-GvpC-H10.H14-His6 | This study |
| pET28a-GvpC-H28.H32-His6 | This study |
| pET28a-GvpC-H10.H14.H17-His6 | This study |
| pET28a-GvpC-H10.H14.H28.H32-His6 | This study |
| pET28a-GvpC-H10.H14.H17.H28.H32-His6 | This study |
| pET28a-GvpC-H6.H10.H14.H17.H20-His6 | This study |
| pET28a-GvpC-H6.H10.H14.H17.H28.H32-His6 | This study |
| pET28a-GvpC-H6.H10.H14.H17.H20.H28.H32-His6 | This study |
| pET28a-GvpC-H6.H10.H14.H17.H20.H24.H28.H32-His6 | This study |
| Bacterial strains | Source |
| *Dolichospermum flos-aquae* CCAP 1403/13F<br>*Escherichia Coli* BL21 (DE3) | CCAP (Culture Collection of Algae and Protozoa)<br>Novagen |

## Acknowledgments

The authors wish to thank Bill Ling, and Mikhail Shapiro for facilitating exploratory in vivo experiments. The authors also thank Shirin Shivaei, Stefan Huber and Arjen Jakobi for their thoughts on sensor designs. The authors also thank Flora Nelissen for support on CCD applications and in vivo experiments. This work was supported by Delft University Fund, the Dutch research council (NWO.STU. 019.021), the Chan Zuckerburg foundations (Dynamic RFA number 2023-321233) and the European Union (Marie-Sklodowska Curie Fellowship MIC-101032769).

## Author contribution

D. Terwiel and D. Maresca conceived and designed the study. D. Terwiel, B. M. Park, and B. Heiles designed and planned experiments. D. Terwiel, B. M. Park, B. Heiles, and E. Muñoz-Ibarra conducted the experiments. D. Terwiel, B. M. Park, B. Heiles, and R. Waasdorp developed software for data collection and analysis. D. Terwiel, B. M. Park, and T. Ara developed bioengineering protocols. D. Maresca and V. Gazzola contributed resources. D. Maresca acquired funding. D. Terwiel, B. M. Park and D. Maresca wrote the manuscript with input from all authors. D. Maresca supervised the research.

## Appendix A Appendix

**Figure A1.**
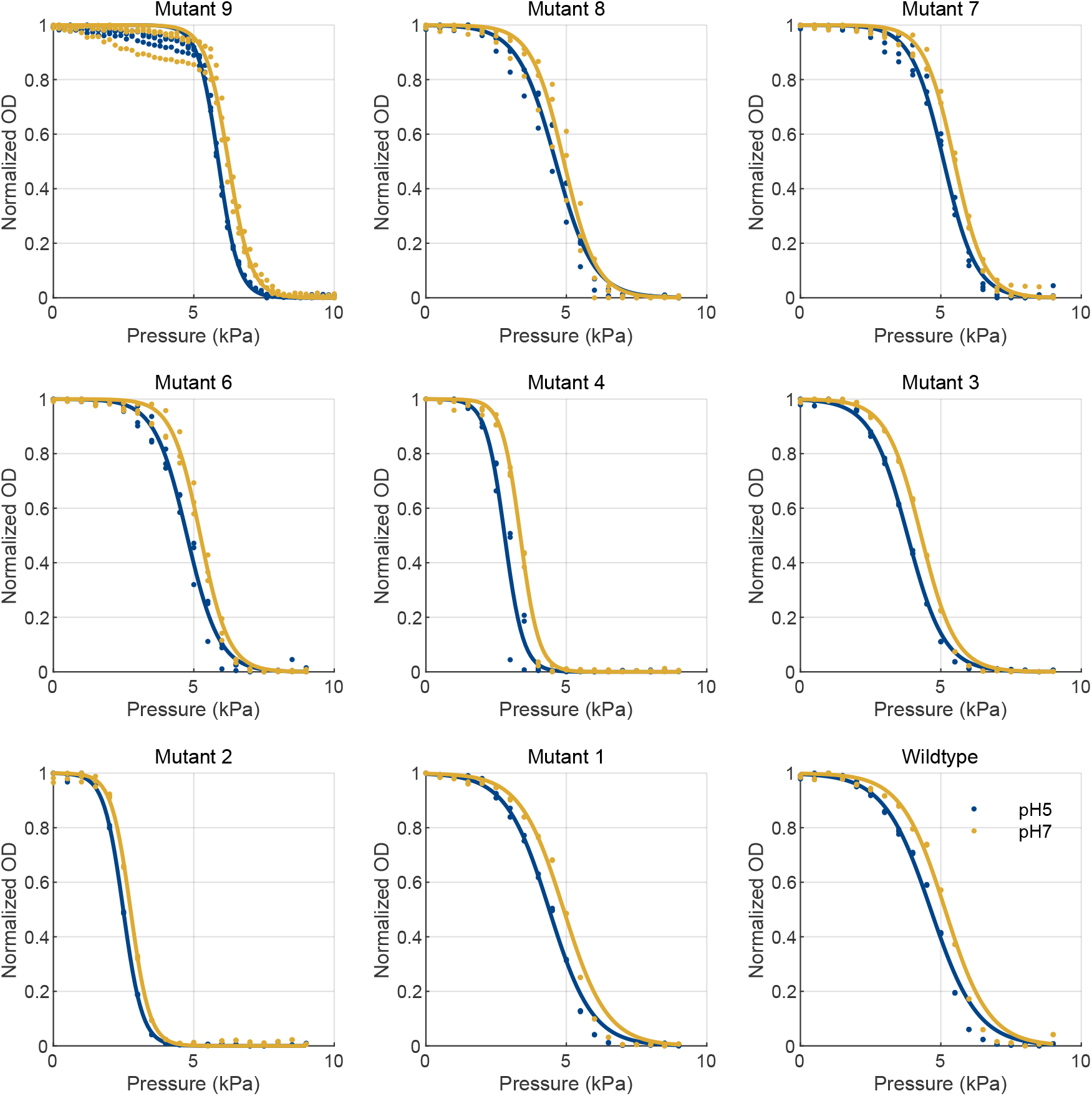
Fitted collapse pressure curves at pH 7 (yellow) and pH 5 (blue) with raw data as scatter plot (n = 3).

**Figure A2.**
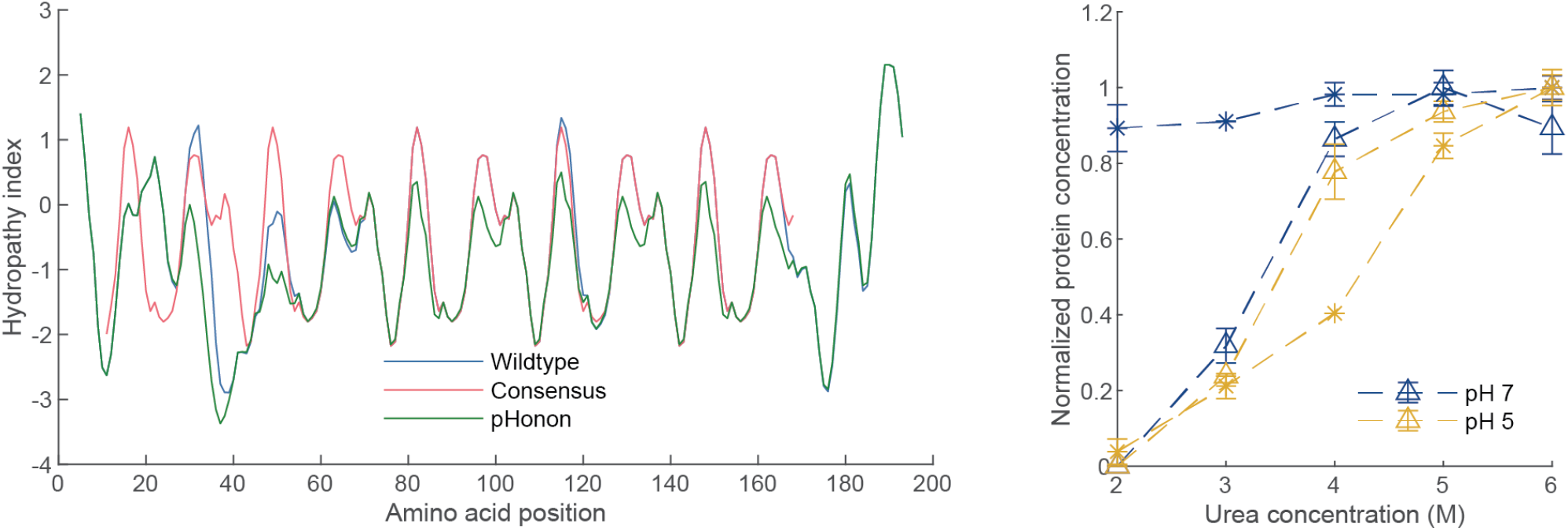
Solubility of GvpC. **a**, kyte-doolittle hydropathy index for wildtype (blue) GvpC, GvpC with a consensus sequence in each of the 5 repeats (red), and pHonon GvpC (blue). pHonon is significantly less hydropathic than wtGvpC. **b**, normalized protein concentration measurements of pHonon (Δ) and GvpC-his (*) when reducing the urea concentration at pH 5 (yellow) and pH 7 (blue). Especially at pH 7, there is almost no precipitation of GvpC-his above 2 M urea, while pHonon GvpC is almost fully precipitated at 2 M urea.

**Figure A3.**
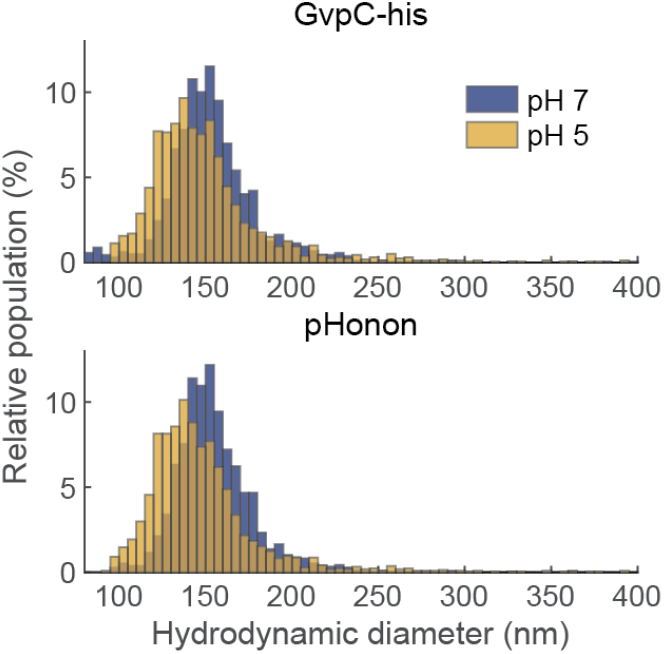
TRPS measurements of GvpC-his GVs and pHonon, at pH 7 (blue) and pH 5 (yellow).

**Figure A4.**
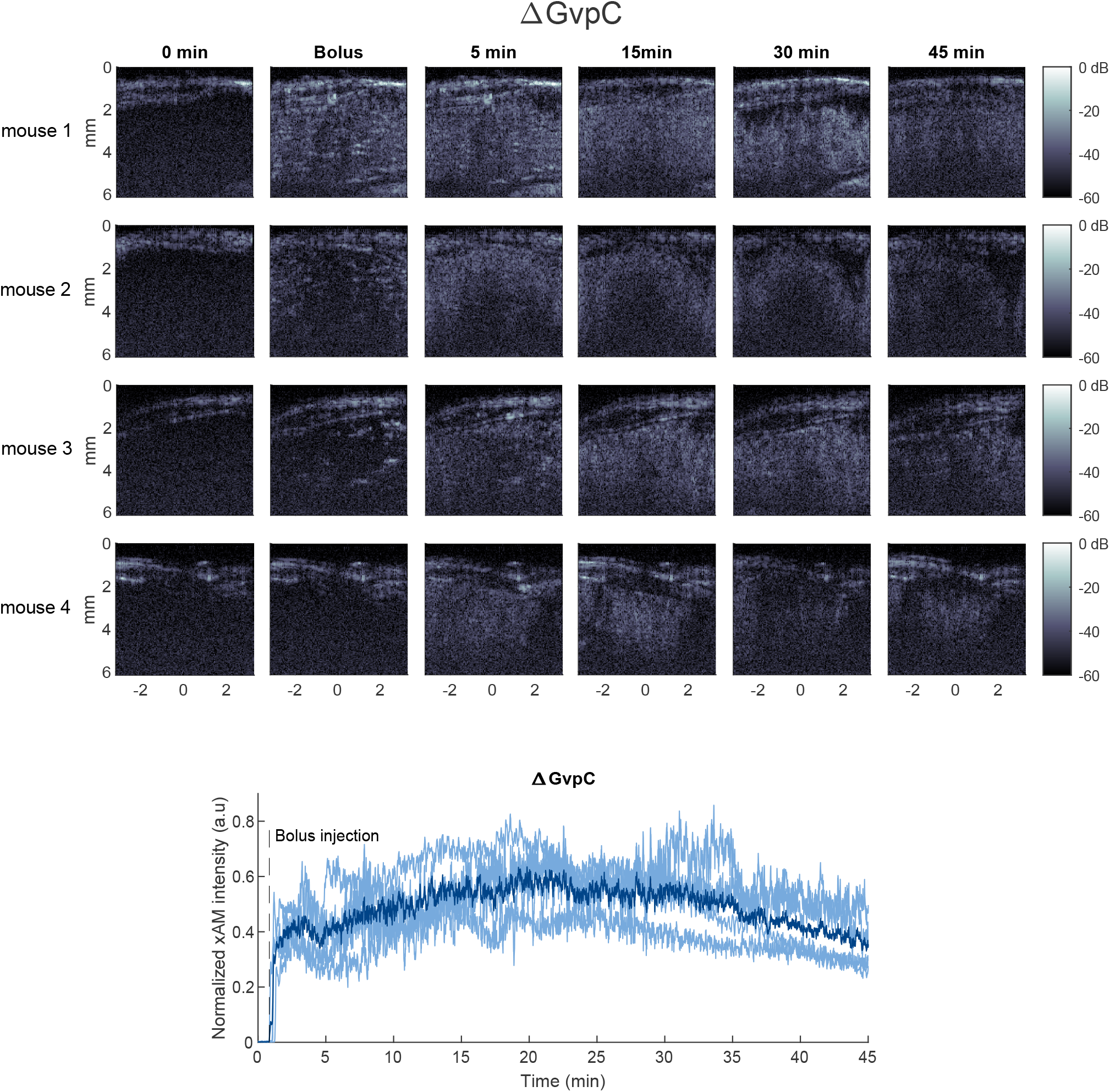
Representative images of liver and time-intensity curves for each separate mouse, with the dark line representing the mean, after injection with ΔGvpC for each of the four mice at different time-points.

**Figure A5.**
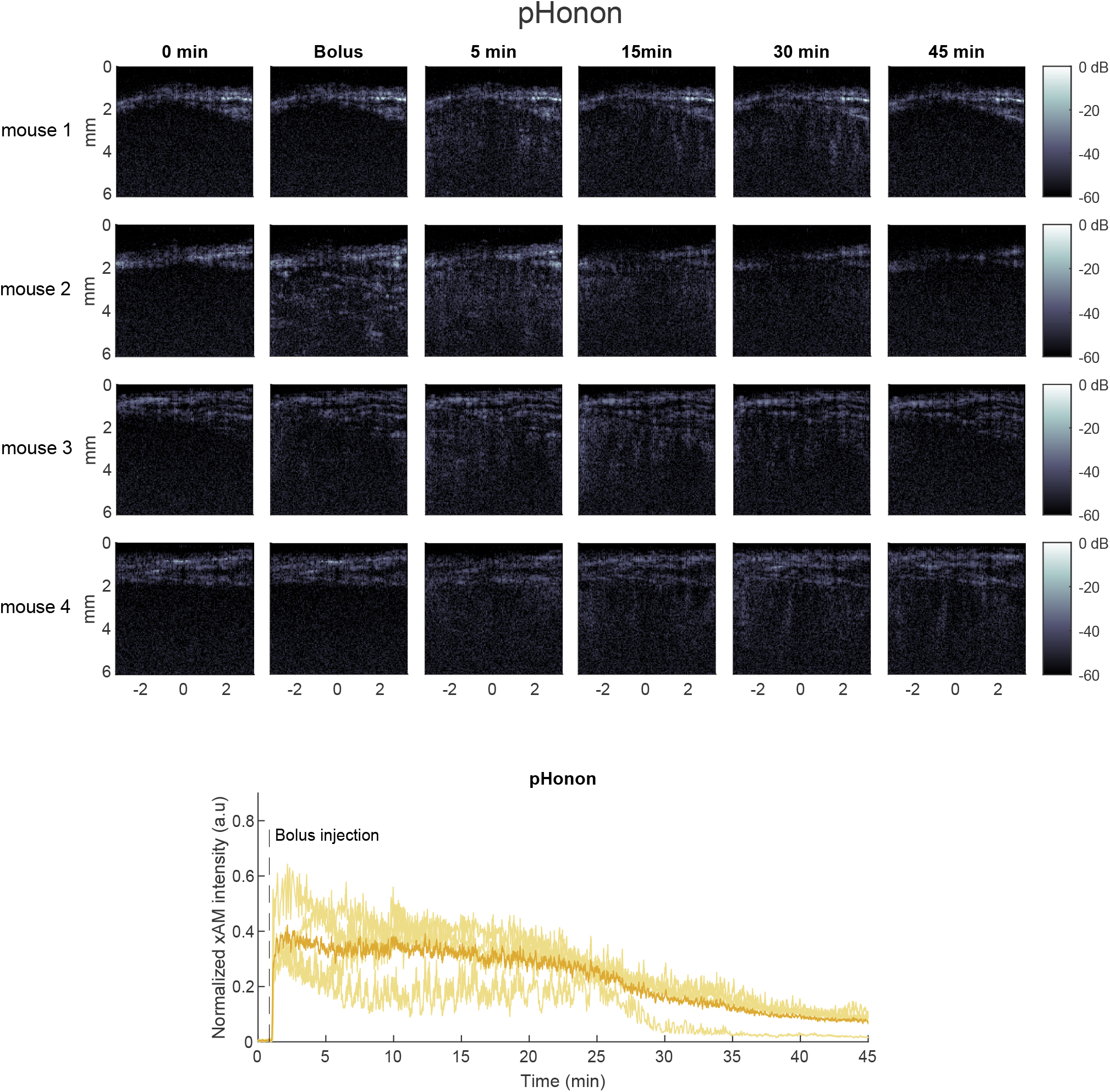
Representative images of liver and time-intensity curves for each separate mouse, with the dark line representing the mean, after injection with pHonon for each of the four mice at different time-points.

